# Age-associated oncocytic transformation correlates with an increased prevalence of small multiple Biondi body inclusions in human choroid plexus epithelial cells

**DOI:** 10.64898/2026.05.27.728253

**Authors:** Caren Yassa, Erfan Zolfaghari, Michael J. Neel, Riley Scanlon, Brett A. Johnson, Edwin S. Monuki

**Author notes:** Correspondence: Edwin S. Monuki.

## Abstract

The choroid plexus epithelial cells (CPECs) at the blood-cerebrospinal fluid (CSF) interface possess an exceptionally high mitochondrial content to support CNS homeostasis. Oncocytic CPECs (O-CPECs), characterized by enlarged and granular eosinophilic cytoplasm composed of excessive abnormal mitochondria, likely contribute to an energetic failure of this energy-demanding tissue. The relationship between O-CPECs and other CPEC pathologies in humans, such as Biondi body (BB) amyloid inclusions, remains poorly defined. In the present study, using H&E-stained sections from 68 postmortem cases, we classified O-CPECs by quantitative size criteria and cytological features, and found an increase in the prevalence of O-CPECs with age after adjusting for sex and tissue source. After excluding two influential control cases, there was evidence for a further increase associated with Alzheimer’s disease. Using antibodies to ATP synthase beta chain to classify O-CPECs, and thioflavin-S to identify BBs, we revealed an increased prevalence of BBs in O-CPECs compared to neighboring non-oncocytic cells. Small multiple BB inclusions were responsible for the increase in O-CPECs, while the prevalence of larger inclusions was decreased in O-CPECs. Together, our data support a clear age-associated oncocytic transformation of CPECs and implicate mitochondrial dysfunction-amyloid interactions.

## INTRODUCTION

Within the brain ventricles, the highly vascularized choroid plexus (ChP) forms a barrier between the blood and the cerebrospinal fluid (CSF), clears the brain’s waste, and produces the CSF (Lehtinen et al, 2013; Lun et al., 2015; Spector et al., 2015; Kaur et al., 2016; Ghersi-Egea et al, 2018). To meet their significant energy demands, ChP epithelial cells (CPECs) contain an exceptionally high mitochondrial volume fraction across species, including rodents and humans, likely rendering them vulnerable to mitochondrial mutations and dysfunction as a tradeoff (Cornford et al., 1997; Xu et al., 2021). With aging and disease, pathologies arise and accumulate in CPECs (Kaur et al., 2016; Reynolds & Putterman, 2025), including, but not limited to, the accumulation of intracellular amyloid inclusions known as Biondi bodies (BBs) (Biondi, 1933; Miklossy et al., 1998; Wen et al., 1999; Ghetti et al., 2024; Neel et al., manuscripts in preparation) and a form of abnormal cellular enlargement known as oncocytic transformation.

Oncocytic transformation occurs when cells become abnormally enlarged and granular-appearing with a strong affinity for eosin stain due to excessive mitochondria (Guaraldi et al. 2011). Oncocytic transformation can arise in many organs and tissues throughout the body, including the adrenal, salivary, pituitary, and parathyroid glands; skeletal muscle, thyroid, kidney, eyes, skin, breast, and multiple brain regions as a compensatory mechanism of dysfunction of mitochondrial enzymes and activity (Hamprel, 1962; Hartwick & Batsakis, 1990; Tallini, 1998; Tanji et al., 2000; Máximo et al., 2002; Guaraldi et al., 2011; Villanueva-Castro et al., 2025).

Oncocytic CPECs (O-CPECs) were initially reported in a patient with Leigh’s disease, a well-known mitochondrial disorder (OMIM# 500017). In this patient, O-CPECs from all four ventricles had enlarged granular cytoplasm and excessive abnormal mitochondria by electron microscopy. Similar O-CPECs were reported in three other patients with Leigh’s disease (Kepes, 1983; Ohama et al., 1988). O-CPECs were also observed in autopsied cases with Kearns-Sayre syndrome (KSS; OMIM# 53000) associated with high levels of mtDNA deletion, reduced expression of mtDNA-encoded cytochrome c oxidase (COX, complexes I/II), and preserved nuclear DNA-encoded mitochondrial respiratory chain components (Tanji et al., 2000). O-CPECs have also been described in two MELAS cases (mitochondrial encephalomyopathy with lactic acidosis and stroke-like episodes; OMIM# 54000) and in a case with multiple mtDNA deletions (Cottrell et al., 2001c).

In addition to primary mitochondrial disorders, enlarged O-CPECs have also been associated with aging as well as neuroinflammatory and neurodegenerative diseases. One study using enzymatic histochemical staining for COX IV and succinate dehydrogenase (SDH, complex II) revealed a distinct subset of granular and enlarged CPECs that were COX-deficient and SDH-positive (COX−/SDH+), which significantly increased with age and Alzheimer’s disease (AD; Cottrell et al., 2001a,b). In another study, laser capture microdissection of COX−/SDH+ CPECs revealed four-fold clonal expansion of mitochondria harboring mtDNA deletions in addition to their enlarged oncocytic phenotype, which accumulated in AD, Parkinson’s disease, and multiple sclerosis (Campbell et al., 2012).

Notably, the multiple mtDNA deletion case (Cottrell et al., 2001c) had a particularly high prevalence of BB inclusions for age (~72%). BBs are lipofuscin-associated amyloid inclusions in CPECs that stain with thioflavin-S (ThS) and have multiple distinctive morphologies and sizes (Biondi, 1933; Oksche & Vaupel-von Harnack, 1969; Oksche & Kirschstein, 1972; Eriksson & Westermark, 1986, 1990a,b; Kitkento, 1986; Neel et al., manuscripts in preparation). BBs can also pierce cell membranes and are spatially clustered within the ChP (Neel et al., manuscripts in preparation). A recent study revealed that transmembrane protein 106B (TMEM106B), which forms amyloid in the aging human brain and is associated with neurodegenerative disease risk (Feng et al., 2021, Perneel et al., 2023), labels BBs (Ghetti et al., 2024). At a population level, BBs abruptly increase in prevalence ~50 years of age and involve the majority of CPECs by ~70 years of age, thereby representing a potential early biomarker of amyloid burden in the brain that may also be associated with AD (Eriksson & Westermark, 1986, 1990a,b; Miklossy et. al, 1998, 1999; Wen et al., 1999; Cottrell et al., 2001c; Neel et al., manuscript in preparation).

In this paper, we characterize O-CPECs in an independent cohort from the UCI Alzheimer’s Disease Research Center (ADRC) and general autopsy service. We report a significant increase in O-CPEC prevalence with age as well as an association with AD. In addition, BB prevalence is increased in O-CPECs, which also have multiple smaller BBs at the expense of having fewer large inclusions.

## RESULTS

### O-CPEC prevalence is positively associated with age

We first identified definite and possible O-CPECs in H&E-stained sections on the basis of their large size, eosinophilic and/or granular cytoplasm, altered and/or displaced nucleus, and marginalized non-nuclear hematoxylin staining (Fig. 1A, Fig. S1A; see formal decision tree in Fig. S2). Focusing first on cell size, histogram analysis revealed a bimodal area distribution corresponding to larger oncocytic cells (median = 316 μm^2^) and smaller non-oncocytes (median = 167 μm^2^) with a significant difference between the two groups (p < 0.0001; Fig. S1A-B). These median areas were larger, but comparable in relative magnitude, to those previously reported (213 μm^2^ and 124 μm^2^ for COX-negative and COX-positive cells, respectively; Cottrell et al., 2001a,b). Based on the histogram analysis, 180 μm^2^ was subsequently used as the minimum threshold for O-CPECs.

**Fig. 1.**
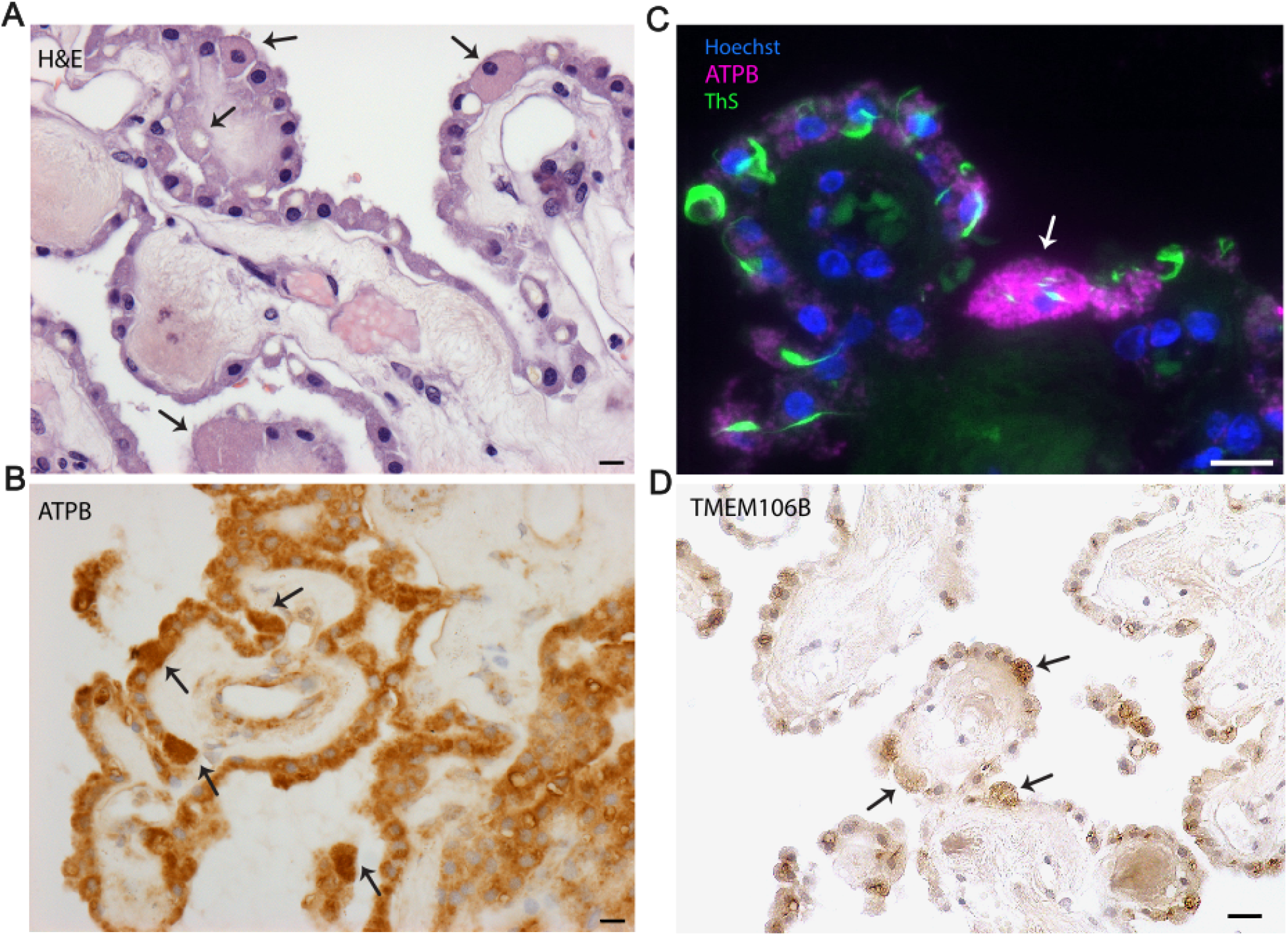
Classification and characterization of O-CPECs. **A**) O-CPECs (**arrows**) appear as enlarged, eosinophilic CPECs in H&E-stained sections. **B**) Due to mitochondrial proliferation, enlarged O-CPECs stain more intensely using antibodies to the beta subunit of the nuclearly encoded mitochondrial protein ATP synthase (ATPB), here visualized by using indirect peroxidase/diaminobenzidine immunohistochemistry. **C**) To analyze Biondi bodies within O-CPECs, we used immunofluorescence for ATPB to highlight O-CPECs together with thioflavin-S (ThS) to label Biondi body amyloid inclusions. **D**) Biondi bodies represent amyloid aggregates of the lysosomal membrane protein TMEM106B, which can be stained using antibodies to the C-terminus after aggressive antigen retrieval. The TMEM106B-positive structures in enlarged O-CPECs typically appear as small multiple inclusions. Scale bars in **A-C** denote 10 *μ*m. Scale bar in **D** denotes 20 *μ*m.

Applying an H&E feature-based decision tree (Fig. S2), we identified O-CPECs in 68 H&E-stained slides chosen from two UCI tissue repositories to represent the human lifespan and to be approximately matched for AD and controls in cases from individuals over 65 years old (Supplemental Table). O-CPECs were sporadic throughout the ChP epithelium with the highest prevalence recorded at 1.9% (Supplemental Table). Across all cases, O-CPEC prevalence increased significantly with age (p < 0.001, R^2^ = 0.652; Fig. 2). Analysis of covariance (ANCOVA), adjusting for age, sex, and tissue source, identified age as the only significant predictor of O-CPEC prevalence (Table 1). There were two subjectively high outliers between 80 and 100 years old (Fig. 1, red symbols) that were determined to be “influential points” based on Cook’s distance analysis (Fig. S3), which identifies the degree of change in the regression model when observations are removed (Cook, 1977). Age remained a significant predictor with these two points removed (Table 1).

**Table 1.** ANCOVA on the percentage of CPECs that are oncocytic.

| | All Cases n=68 | | Cases $\geq 65$ n=40 | |
| --- | --- | --- | --- | --- |
|  | w/ Influential Points | w/o Influential Points | w/ Influential Points | w/o Influential Points |
| Age | <0.001* | <0.001* | <0.001* | 0.001* |
| Sex | 0.278 | 0.290 | 0.309 | 0.421 |
| AD | N/A | N/A | 0.800 | 0.022* |
| Tissue Source | 0.535 | 0.939 | 0.830 | 0.061 |

**Fig. 2.**
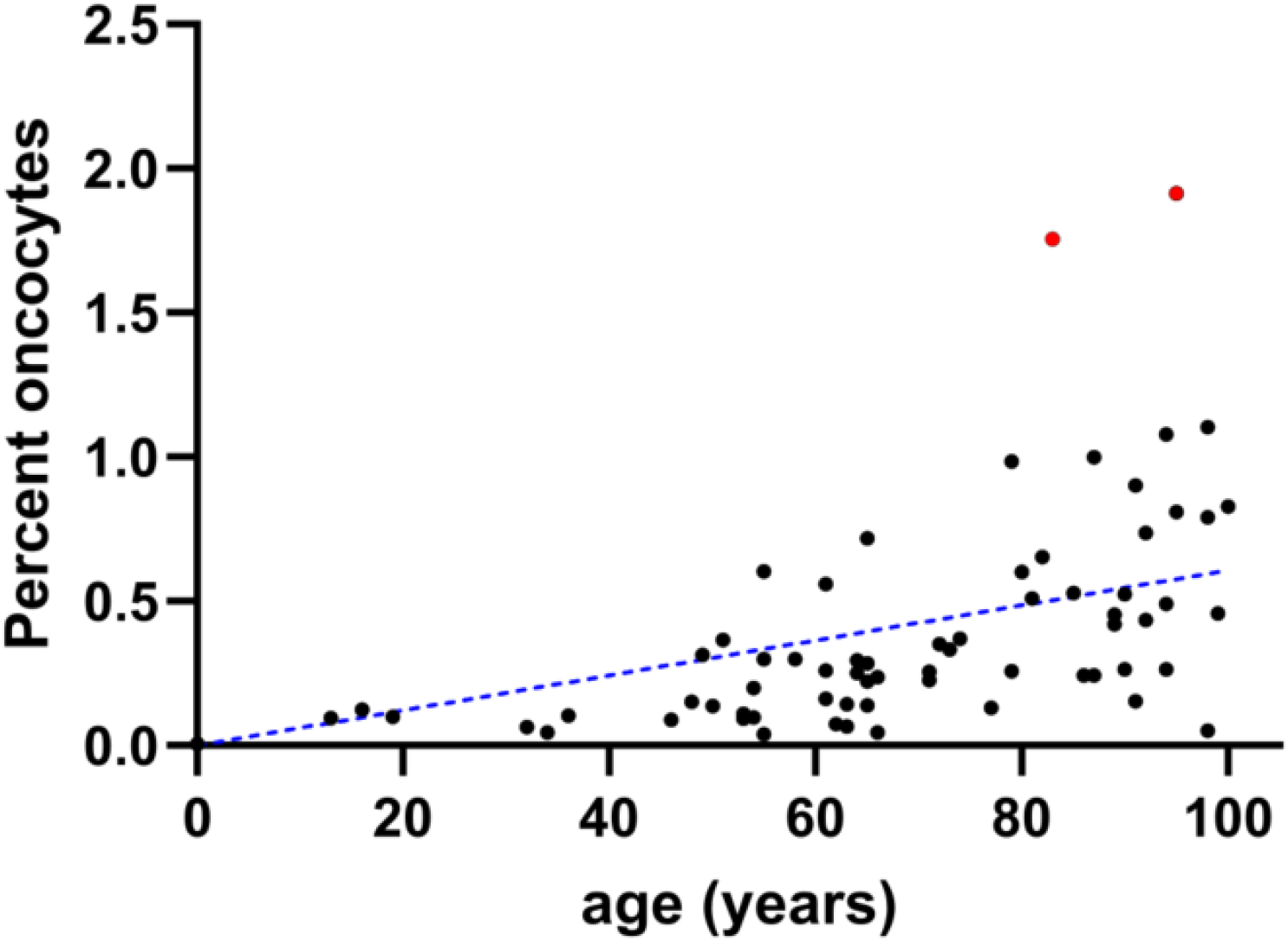
O-CPECs accumulate with age. H&E-stained sections from 68 individuals were assessed for the percentage of CPECs with oncocytic characteristics. The line is a result of simple linear regression. Red plot symbols denote influential data points in Cook’s distance analysis that were treated as outliers in some subsequent statistical treatments.

### O-CPEC prevalence is positively associated with Alzheimer’s disease

Analyses were repeated for cases >65 years of age for which AD status was available (n=40 of 68, 17 AD, 23 non-AD). O-CPEC prevalence remained significantly increased with age in this >65 years-old cohort (p<0.001), whereas no significant associations were observed with AD, sex, or tissue source. However, excluding the two influential points identified by Cook’s distance (which were both non-AD cases) revealed a significantly increased prevalence of O-CPECs in AD cases (p=0.022; Fig. S3). Sex and tissue source remained unassociated (Table 1).

### O-CPECs and BBs can be co-visualized using anti-ATPB antibody and ThS

A recent study reported that BBs were aggregates of TMEM106B (Ghetti et al., 2024). Indirect TMEM106B immunoperoxidase staining with 3,3′-diaminobenzidine (DAB) revealed BB-positive inclusions in enlarged CPECs and gave the subjective impression of generally smaller BB morphologies in enlarged compared to normal-sized CPECs (Fig. 1D). To co-identify O-CPECs and BBs more clearly, we tested an antibody for ATPB, a subunit of the ATP synthase complex that is primarily encoded by nuclear DNA (Lai et al., 2023). Intensity of DAB staining for ATPB distinguished O-CPECs from adjacent non-oncocytic cells (Fig. 1B). We then developed a co-staining procedure using ThS followed by indirect ATPB immunofluorescence, which allowed for BB detection and the straightforward identification of O-CPECs (Fig. 1C).

### O-CPECs have an increased prevalence of BBs and more multiple small inclusions, but fewer large inclusions

We specifically examined the number and greatest dimensions of BBs in O-CPECs and non-oncocytic CPECs, which were classified into six categories:ThS-negative, small few inclusions (SFI), large few inclusions (LFI), small multiple inclusions (SMI), large multiple inclusions (LMI), and small plus large multiple inclusions (SMI+LMI) (Fig. 3C; see Methods for details). (There were no CPECs within the analyzed set that were either SMI+LFI or SFI+LMI.) Overall, BB prevalence was increased in O-CPECs compared to non-oncocytic CPECs (paired t-test, p = 0.0028). In addition, SMI were significantly increased (p <0.0001) and LFI were significantly decreased (p<0.001) in O-CPECs compared to non-oncocytic CPECs, while the other categories were fractionally small in both groups (Fig. 3A-B, Fig. 4A-C; Table 2).

**Table 2.** Paired t-test analyses of differences in morphological categories (O-CPECs minus control CPECS). Asterisks in the rightmost column represent p<0.05 after Bonferroni correction for six comparisons.

| Category | Mean of Differences +/- SEM | p-value | Bonferroni Corrected p-value |
| --- | --- | --- | --- |
| ThS Positive | 13.58 +/- 3.01 | 0.0028* | * |
| Small Few Inclusions | 2.59 +/- 2.11 | 0.2592 | ns |
| Small Multiple Inclusions | 46.54 +/- 4.92 | <0.0001 * | * |
| Large Few Inclusions | -41.30 +/- 3.10 | <0.0001* | * |
| Large Multiple Inclusions | 2.95 +/- 1.42 | 0.0755 | ns |
| Large+Small Multiple Inclusions | 1.85 +/- 0.97 | 0.0977 | ns |

**Fig. 3.**
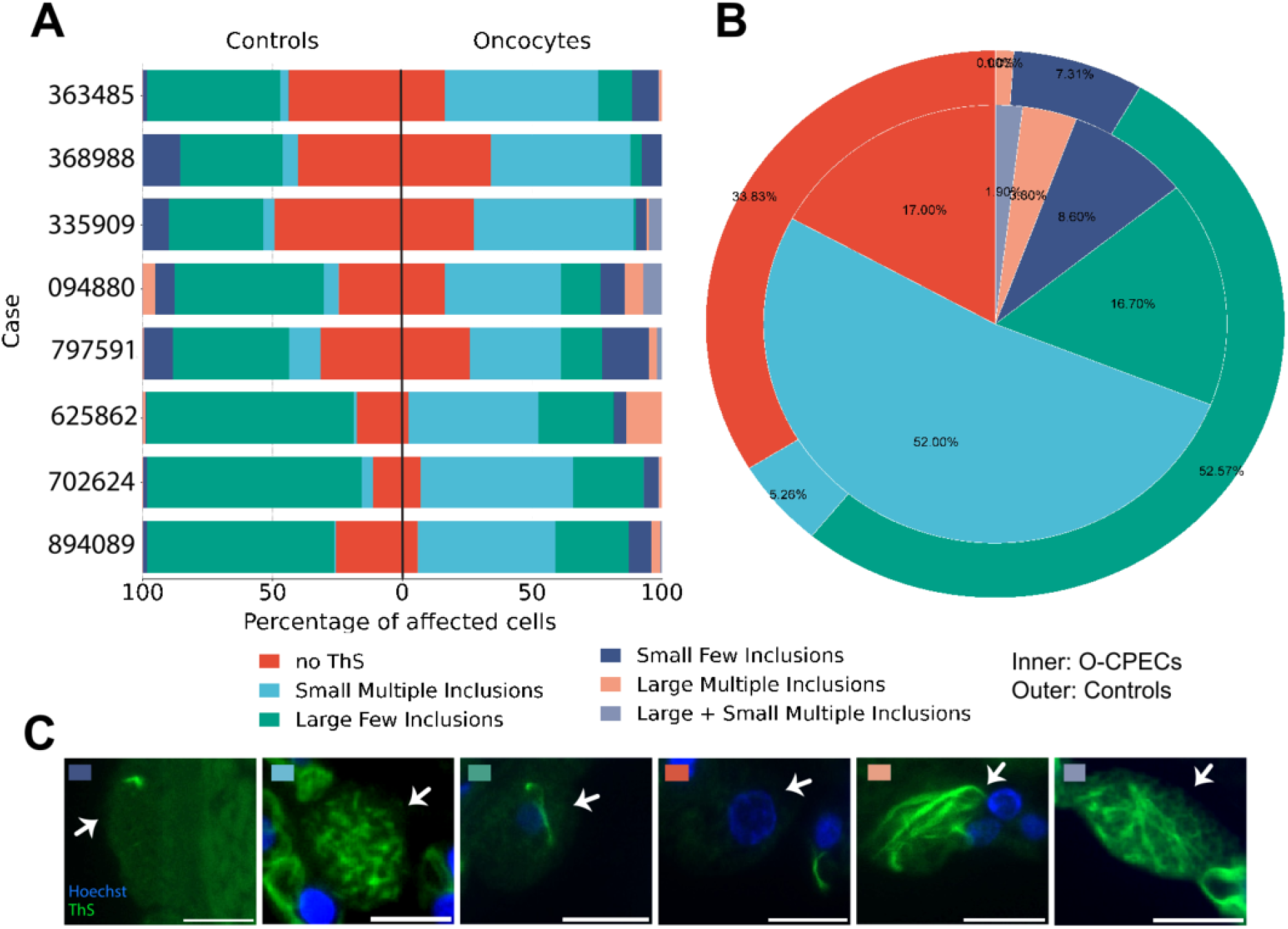
O-CPECs differ from control CPECs in the proportions of BBs with different morphological characteristics. **A**) Bidirectional plots displaying fractions of O-CPECs and control CPECs possessing six distinct morphological categories for each of the eight postmortem cases analyzed. **B**) Concentric pie chart showing the average percentages of O-CPECs (outer ring) and control CPECs (inner circle) across the eight postmortem cases. **C**) Examples of each morphological category. Scale bars are 10 *μ*m.

**Fig. 4.**
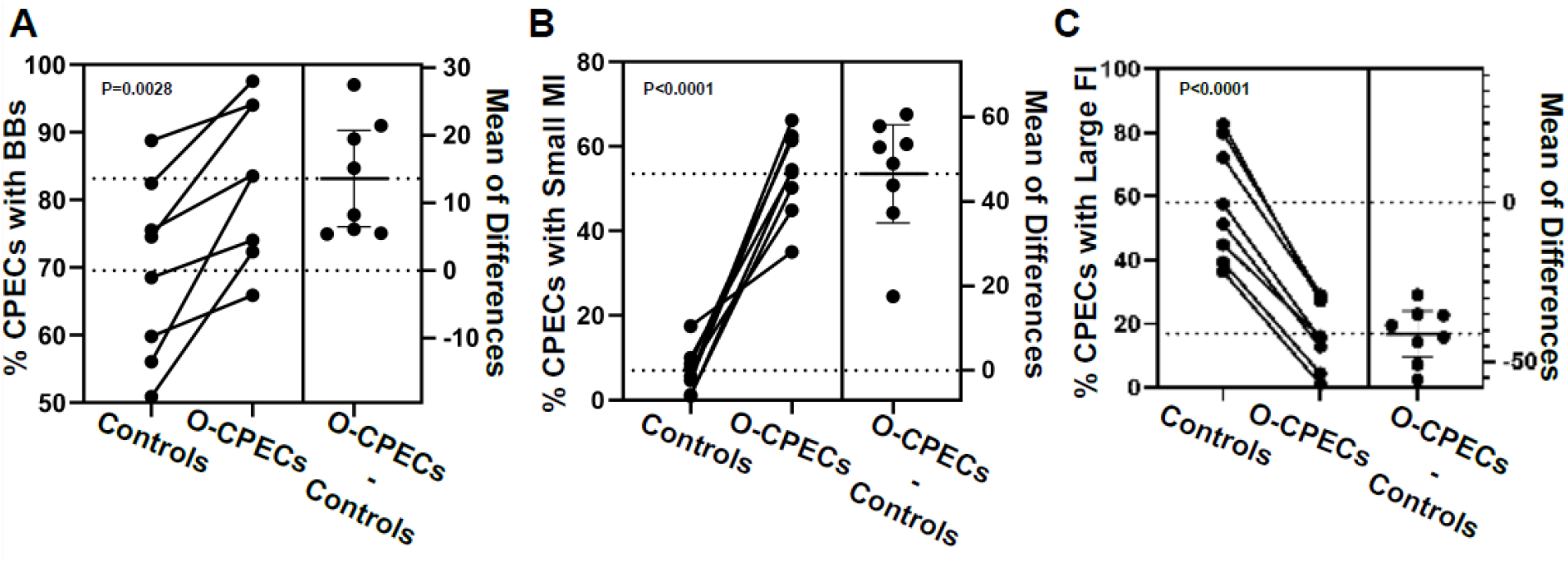
Plots of the paired analyses showing significant differences between O-CPECs and control CPECs in overall percentages of Biondi bodies (**A**), percentages of cells containing small multiple inclusions (**B**), and percentages of cells containing few, large inclusions (**C**).

## DISCUSSION

In this study, we developed a decision tree for O-CPECs based on histologic features in conventional H&E stains, then applied this decision tree to a postmortem cohort of 68 cases across the human lifespan. Using whole slide images and manual annotations, we confirmed the positive association of O-CPEC prevalence with age seen previously (Cottrell et al., 2000, 2001a,b). After formally determining and removing two influential cases (“outliers”) based on Cook’s distance analysis (Cook, 1977), a previous association of O-CPEC prevalence with AD (Campbell et al., 2012) was also confirmed in our >65 years of age cohort. Using a new method to co-visualize O-CPECs with BBs, we then investigated interactions between these two cell-intrinsic pathologies. Overall, O-CPECs displayed an increased prevalence of BB inclusions. In addition, after categorizing these inclusions based on number and size, we found that O-CPECs were markedly enriched for multiple small inclusions at the expense of fewer large inclusions.

### Likely underestimation of O-CPEC prevalence

O-CPEC prevalence values reported here and in other cited end-point studies are relatively low (e.g., less than 2% of CPECs in the current study). It is therefore important to note that the reported prevalence values almost certainly underestimate O-CPEC transformation over time. Epithelial barriers are well known to regularly shed unwanted or undesirable cells via a highly conserved process known as cell extrusion (Villars and Levayer, 2022; Grata and Levayer, 2025), and multiple observations suggest CPEC extrusion from human ChP epithelium. For example, in histological preparations, occasional O-CPECs (and non-oncocytic CPECs) detached from the underlying basement membrane are commonly observed (data not shown), and benign CPECs are rare, but well known to cytologists and neurologists as constituents of CSF taps. Thus, O-CPEC extrusion from ChP epithelium into ventricular CSF is highly likely. If so, extruded O-CPECs would no longer be visible histologically, and the prevalence values reported here would represent O-CPEC *incidence* values over some unit of time. Coupled with the relatively low to absent turnover of CPECs in adult primates and humans (Kaplan, 1980; Korzhevskii, 2000), O-CPEC transformation over the human lifespan is likely much higher than reported here, and accordingly, more easily envisioned as a major contributor to the physical and functional ChP atrophy that occurs with aging and neurodegeneration.

### O-CPEC interactions with BBs

Our finding of increased BB prevalence in O-CPECs is consistent with the increased BBs for age observed in a multiple mtDNA deletion case (Cottrell et al., 2001c). Directionality of this association between O-CPECs and BBs remains to be resolved; O-CPEC transformation may predispose to BB formation, and/or vice versa. For the Cottrell et al. case, the mitochondrial defects predisposing to BB formation seems more likely. That being said, the directionality of O-CPEC transformation itself is clear, i.e., non-oncocytic CPECs, which start with fewer large inclusions, transform into O-CPECs, which end up with multiple small inclusions. Thus, most likely, O-CPEC transformation results in larger BB inclusions becoming multiple smaller ones. A likely explanation for this directional relationship would be physical fragmentation of large BBs by the overproliferating mitochondria of O-CPECs, which are known to have untoward physical impacts on cells (hypertrophy) and cellular organelles (e.g., indentation, margination). Such BB fragmentation would also be consistent with the irregular shapes, sizes, and contours of the multiple ThS-stained inclusions seen in many O-CPECs (data not shown). Another recent study from our laboratory (Neel et al., manuscript in preparation) found a negative association between AD and multiple small BBs in CPECs. On the surface, these findings seem inconsistent, but the Neel et al. study assessed *all* CPECs rather than the relatively rare O-CPECs studied here. Thus, the cell population-based observations and associations in these two studies are largely independent.

### Mitochondrial dysfunction associated with O-CPECs, amyloids, aging, and neurodegenerative diseases

Previous studies have linked mitochondrial dysfunction to O-CPECs and oncocytic transformation in general, while others, including the current study, have linked O-CPECs to aging and AD. Independent of O-CPECs and other oncocytic cell types, there is also a sizable literature linking mitochondrial dysfunction to brain aging, AD, and other neurodegenerations. Within this rich network of mitochondrial associations, directionality of several edges remains unclear, although O-CPEC transformation due to mitochondrial dysfunction is intuitively direct, i.e., mutant mitochondria cause energetic and metabolic deficits in a cell, which result in compensatory, yet ill-fated, mitochondrial proliferation that overwhelms organelle and other control mechanisms within the same cell (Hamprel, 1962; Askew et al., 1971; Sun et al., 1975; Hartwick & Batsakis, 1990; Tallini, 1998; Linnane et al., 1989; Gasparre et al., 2011). Interestingly, several studies imply bidirectional relationships between mitochondrial dysfunction/mtDNA mutations and the accumulation of other amyloid aggregates in the brain (Reddy et al., 2008; Yao et al., 2009; Kukreja et al., 2014; D’Alessandro et al., 2025), which may also be the case for O-CPECs and BBs. The existing literature centers on particularly energy-demanding cells (e.g., neurons) and organs (e.g., human brain) with energy demands and deficits standing out as unifying themes. Findings from this study add the energy-demanding ChP and its CPECs, as well as a new set of CPEC-intrinsic (O-CPEC-BB) interactions to this literature, which may be amenable to more direct and intuitive dissection of these complex mitochondrial and energy-related relationships.

## METHODS

### Human Tissue

Postmortem human ChP samples (glomus) from the atrium (trigone) of the lateral ventricle were obtained through the University of California, Irvine Medical Center (UCIMC) autopsy service and the Alzheimer’s Disease Research Center (ADRC) tissue repositories. Each sample was fixed in either 10% formalin (UCIMC) or 4% paraformaldehyde (ADRC), and embedded in paraffin. Blocks from 68 cases were sectioned at 5-µm thickness and either delivered to us on unstained slides or stained using H&E by the UCI Experimental Tissue Resource.

### Immunostaining

#### ATPB vs ThS

For the 8 cases chosen for the O-CPECs and BBs study, paraffin-embedded 5 µm sections were heated at 60°C for 20 mins, deparaffinized in xylene, rehydrated in a descending alcohol dilution series, treated with 0.5% (w/v) thioflavin-S (ThS, Sigma) in 50% ETOH for 10 mins, followed by 2x 5-min washes in 50% ETOH, and 3x 5-min washes in Tris-buffered saline (TBS). All the steps after the ThS treatment were performed in the dark by covering all the containers used with aluminum foil. Sodium citrate pH 6.0 was used for antigen retrieval in a vegetable steamer for 20 mins, followed by 2x 5-min washes in MilliQ water and 2x 5-min washes in TBS at RT. Hydrophobic Pap-Pen barriers (Research Products International) were drawn around the tissue section and left to dry for a few minutes at RT. The sections were then incubated in blocking solution (5% donkey serum and 0.3% Triton X–100 in TBS) at RT for 1 hr, followed by incubation in primary mouse monoclonal antibody against ATPB (Abcam, ab14730) solution (primary antibody at 1:50 dilution in 1% donkey serum and 0.3% triton X-100 in TBS) at 4°C overnight. The following day, the primary antibody solution was removed, followed by 3x 5-min washes with TBS, incubated in Alexa-647 donkey anti-mouse secondary antibody solution (secondary antibody at 1:200 dilution in 1% donkey serum and 0.3% Triton X-100 in TBS) at RT for 1 hr, then washed 3x 5 mins with TBS. The sections were incubated in Hoechst 33342 solution (2ug/mL in PBS) (Invitrogen) at RT for 5 mins, followed by 3x 5-min washes in TBS. The slides were mounted with 5-6 drops of Fluoromount G (Southern Biotech) and coverslipped.

#### TMEM106B DAB

Following antigen retrieval in 98% formic acid followed by 10 mM sodium citrate, pH 6 heat treatment (Perneel et al., 2023), 5-micron FFPE sections were stained using a 1/6000 dilution of anti-TMEM106B (239-250) (Cosmo TIP-TM-P01) with detection by a Vectastain elite peroxidase-DAB kit, followed by a light hematoxylin counterstain.

### Image analysis and quantification

#### H&E images quantification

Whole slide images were scanned using an Aperio Versa 2 scanner in the ADRC neuropathology core. The whole-slide scanned images were opened in ImageScope software (v12.4, Leica Biosystems) and viewed at 40X magnification. An estimate of the total number of CPECs was determined using a previously described and validated deep-learning model (Espericueta et al., 2026). Analysis of O-CPECs typically involved the entire whole slide image, although in some sections that were very large, analysis was terminated at the 60,000th CPEC detected by the deep learning model.

O-CPECs were manually quantified after specialized training from a project scientist (BAJ) who was trained by a board-certified neuropathologist (ESM), followed by multiple feedback sessions to refine annotation accuracy. Taking advantage of previously characterized oncocytic features and sizes, a few cases were first analyzed to identify definite, possible, and non-oncocytes defined by a certified neuropathologist (ESM). Analysis of the two-dimensional area of the three categories revealed a bimodal distribution between definite oncocytes and non-oncocytes, while possible oncocytes overlapped at the antimode (Fig. S1). A 2-D area cutoff (180 μm^2^) was determined at the border of increased frequency of probable and definite oncocytes. A considerable number of non-oncocytes also possessed cross-sectional areas above 180 μm^2^ (Fig. S1), which necessitated further steps to classify the CPECs. Based on literature descriptions of oncocytes throughout the body (Guaraldi et al., 2011), features most frequently present in the oncocytes identified by the neuropathologist, and features most consistently recognized across different annotators, a decision tree was generated to standardize classification of oncocytes across different annotators (Fig. S2). The highest priority feature was size, followed by eosinophilicity, nuclear changes, and finally, granular cytoplasm, nuclear displacement, and marginalized non-nuclear hematoxylin staining (likely endoplasmic reticulum). Very large O-CPECS confidently identified by the neuropathologist did not always express many of the other features, so the decision tree required fewer features with increasing size. Annotators, who were unaware of ages, sexes, and AD status, followed the decision tree criteria for all 68 H&E-stained cases.

#### IHC images quantification

Blinded to the ThS channel, O-CPECs were characterized by a brighter ATPB signal relative to neighboring cells and an area greater than 180 μm^2^ using the tracer tool on the software. Three adjacent non-oncocytic CPECs on each side of the marked oncocytic cell were annotated within the same monolayer, and all cells were labeled using the counter tool. BBs were then classified into six different categories based on the BB morphologies in the cells marked with the counter tool, blinded to the ATPB channel. The six different BB morphology categories included: no ThS detection, small few inclusions (<3 + <4µm in greatest dimension), large few inclusions (<3 + >4 µm in greatest dimension), small multiple inclusions (≥ 3 + <4µm in greatest dimension), large multiple inclusions (≥ 3 + >4µm in greatest dimension), and small plus large multiple inclusions (≥ 3 of <4 µm in greatest dimension + ≥ 3 of >4 µm in greatest dimension).BB lengths were measured using the ruler tool to determine the reference length of 4µm. The length was transferred from the screen to a transparent sheet and doubled to serve as a scale for measurement in the magnifier window on the software (~80X magnification: 2X magnifier digital zoom over 40X optical scan).

### Statistical analysis

GraphPad Prism 9 software and SRplot were used for graphing and statistical analysis. Analysis of covariance (ANCOVA) was conducted using SPSS version 28 software with robust error to account for heteroskedasticity in the data. The specific tests used are described in the figure legends. Ggplot2 package (version 3.5.2) in R was used for Cook’s distance analysis.

## ACKNOWLEDGEMENTS

We are deeply indebted to the tissue donors and their families for enabling this research. We thank UCI neuropathologists and technicians at the Experimental Tissue Resource and UCI ADRC Neuropathology Core for their assistance in tissue procurement, tissue processing, and slide scanning. This work was supported by the following NIH awards: P50AG016573 and P30AG066519 (UCI ADRC), T32AG073088 and T32AG000096 (M.J.N.), and R21MH109036 and R21AG064640 (E.S.M.).

## FIGURES

**Fig. S1.**
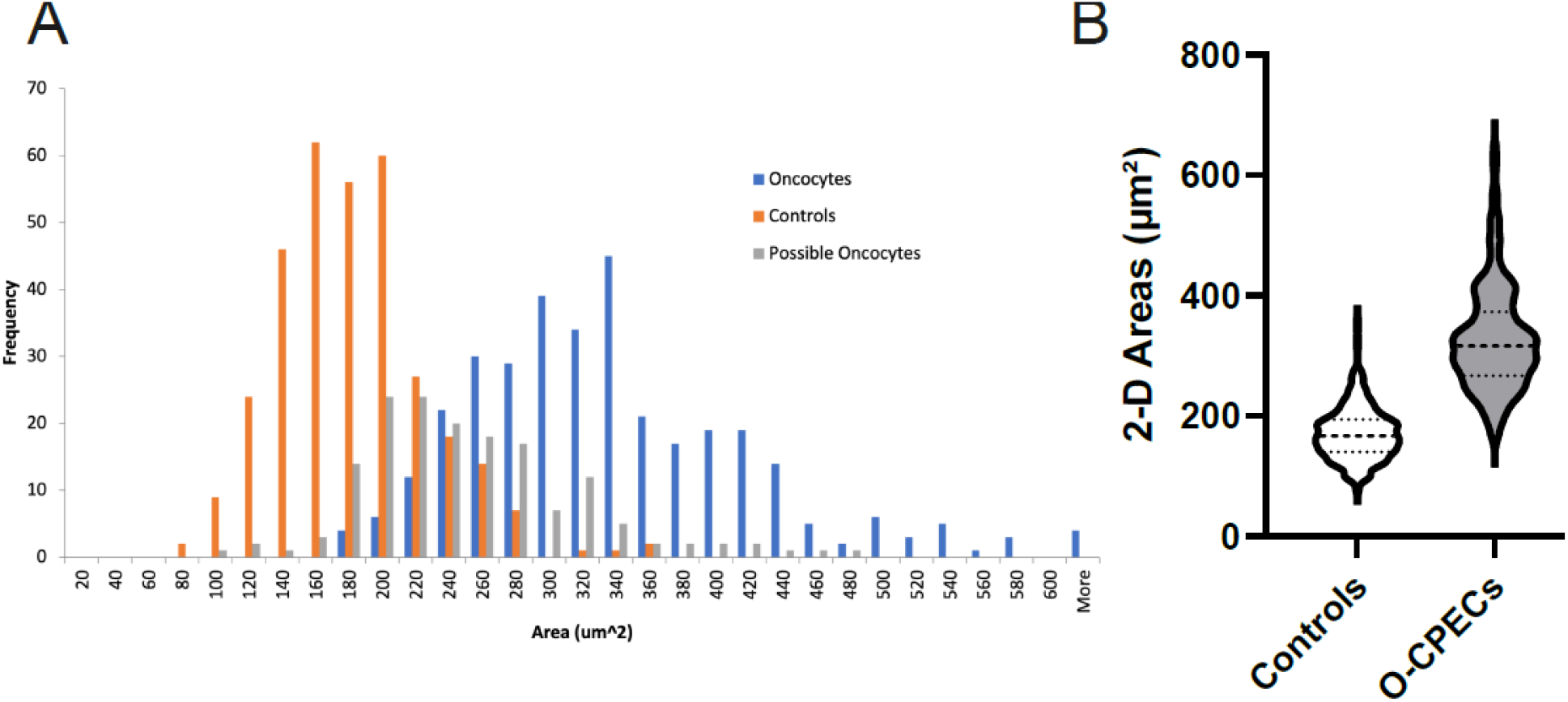
The cross-sectional areas (5 *μ*m sections) of potential O-CPECs and a collection of control CPECs were measured after being classified by a board-certified pathologist (ESM) according to their histological features. **A**) Frequency of observed areas in each of three categories. **B**) Violin plots of O-CPECs and reliably defined control CPECs. A Mann-Whitney U test indicated that sizes were significantly different between the histologically defined categories (p<0.0001).

**Fig. S2.**
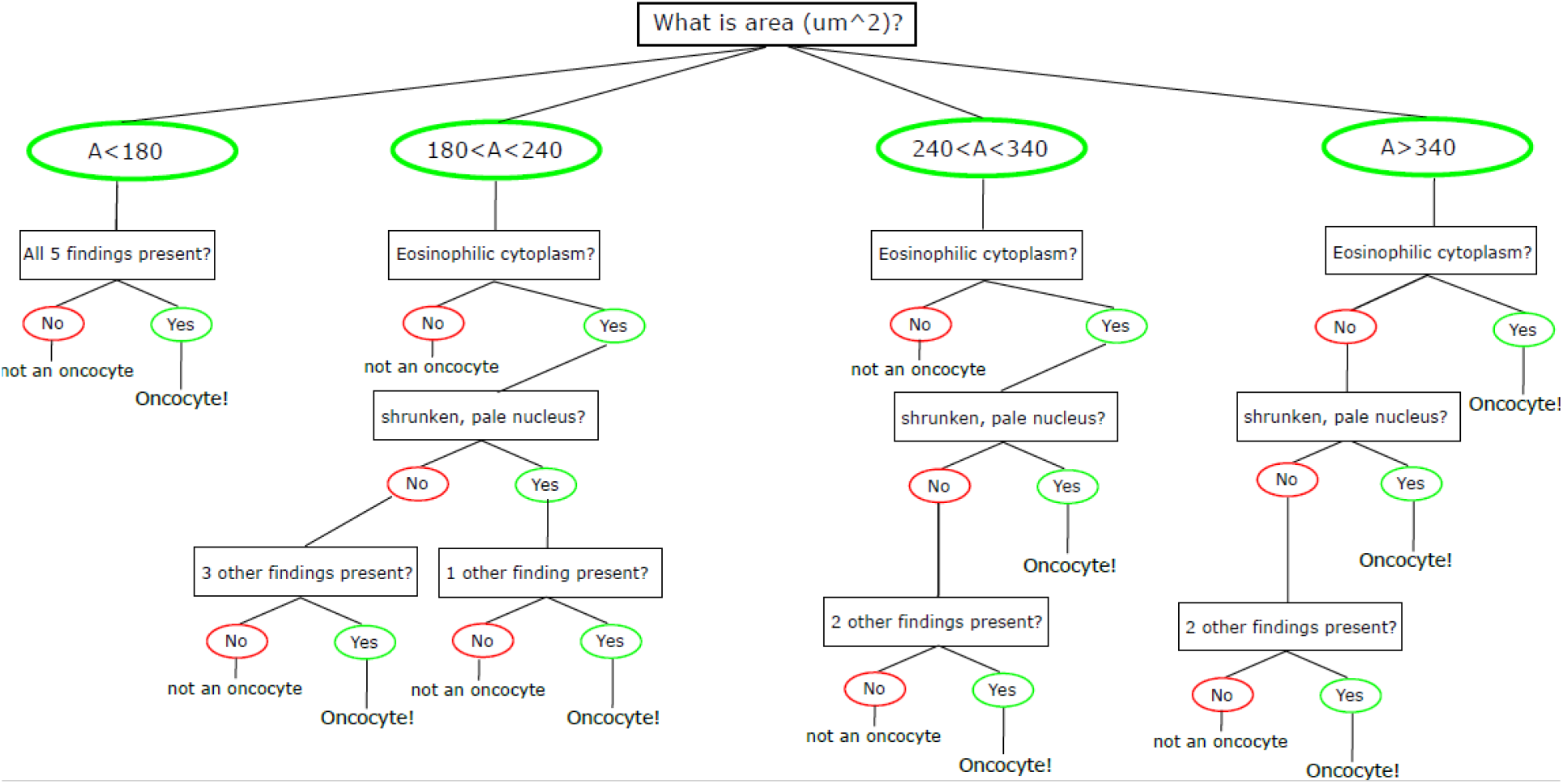
A decision tree was used to standardize classification of O-CPECs in H&E-stained sections. In addition to the characteristics explicitly stated, the remaining three determinants included cytoplasmic granularity, nuclear displacement, and marginalized hematoxylin.

**Fig. S3.**
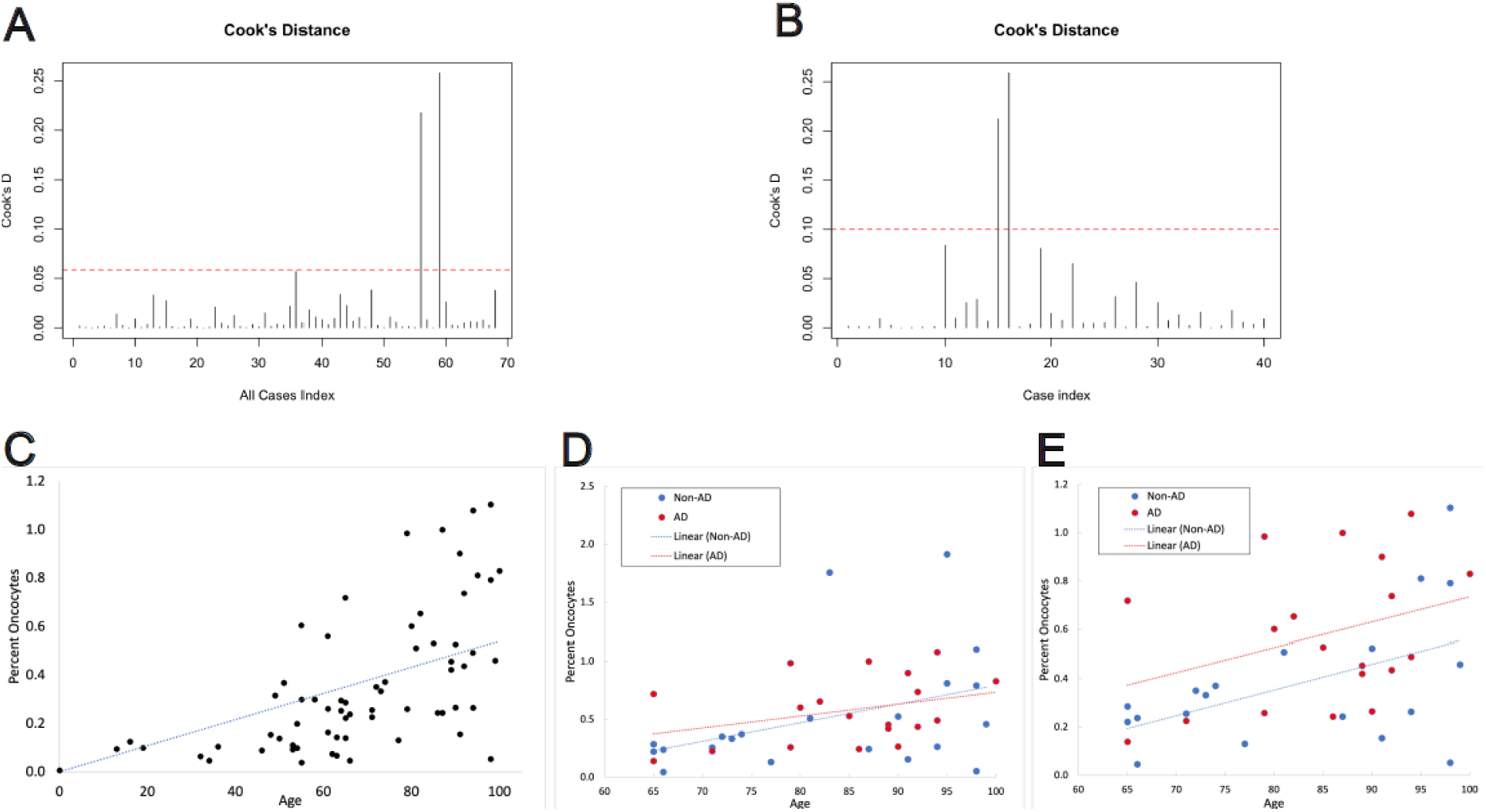
Cook’s distance analysis identified “influential” outlying data points that affected the results of statistical analyses. **A**,**B**) Plots of Cook’s distance for points across the lifespan (**A**) and points ≥ 65 years old for analyzing the effects of Alzheimer’s disease (**B**). Two data points exceeded the threshold values (4/n) and were identified as influential. **C**) A plot of O-CPEC prevalence with the outlying data points removed, which can be compared to Fig.2. **D, E**) Plots of data for individuals ≥ 65 years old with (**D**) and without (**E**) the influential datapoints identified by Cook’s distance analysis. Lines are the result of simple linear regression.

